# Lymphatic type-1 interferon responses are critical for control of systemic reovirus dissemination

**DOI:** 10.1101/2020.07.11.198267

**Authors:** Matthew B. Phillips, Marcelle Dina Zita, Morgan A. Howells, Tiffany Weinkopff, Karl W. Boehme

**Author notes:** Correspondence to Karl W. Boehme, Mailing address: 4301 W. Markham Dr. #511, Little Rock, AR 72205.

## Abstract

Mammalian orthoreovirus (reovirus) spreads from the site of infection to every organ system in the body via the blood. However, mechanisms that underlie reovirus hematogenous spread remain undefined. Non-structural protein σ1s is a critical determinant of reovirus bloodstream dissemination that is required for efficient viral replication in many types of cultured cells. Here, we used the specificity of the σ1s protein for promoting hematogenous spread as a platform to uncover a role for lymphatic type-1 interferon (IFN-1) responses in limiting reovirus systemic dissemination. We found that replication of a σ1s-deficient reovirus was restored to wild-type levels in cells with defective type-1 interferon a-receptor (IFNAR1) signaling. *In vivo*, reovirus spreads systemically following oral inoculation of neonatal mice, whereas the σ1s-null virus remains localized to the intestine. We found that σ1s enables reovirus spread in the presence of a functional IFN-1 response, as the σ1s-deficient reovirus disseminated comparably to wild-type virus in IFNAR1^-/-^ mice. Lymphatics are hypothesized to mediate reovirus spread from the intestine to the bloodstream. IFNAR1 deletion from cells expressing lymphatic vessel endothelium receptor-1 (LYVE-1), a marker for lymphatic endothelial cells, enabled the σ1s-deficient reovirus to disseminate systemically. Together, our findings indicate that IFN-1 responses in lymphatics limit reovirus dissemination. Our data further suggest that the lymphatics are an important conduit for reovirus hematogenous spread.

**IMPORTANCE:** Type-1 interferon (IFN-1) responses are a critical component of the host response to viral infections. However, the contribution of specific cell and tissue types to control of viral infections is not known. Here, we found that reoviruses lacking nonstructural protein were more sensitive to IFN-1 responses than wild-type reovirus, indicating that σ1s is important for reovirus resistance to IFN-1 responses. The σ1s protein also is a key determinant of reovirus systemic spread. We used tissue-specific IFNAR1 deletion in combination with the IFN-1-sensitive σ1s-null reovirus as a tool to identify a role for lymphatics in reovirus dissemination. We used Cre-lox technology to delete type-1 interferon a-receptor (IFNAR1) in lymphatic cells and found that the IFN-1-sensitive σ1s-deficient reovirus disseminated in mice with lymphatic endothelial cells-specific deletion of IFNAR1. Together, our results indicate that IFN-1 responses in lymphatics are critical for controlling reovirus systemic spread.

## INTRODUCTION

Systemic dissemination is a fundamental step in viral pathogenesis. To spread within the host, viruses must replicate in multiple cell and tissue types and overcome a variety of physical and physiological barriers, including host antiviral defenses. Mammalian orthoreovirus (reovirus) is a member of the *Reoviridae*, a family of non-enveloped, dsRNA viruses that infects via respiratory or enteric routes and subsequently traffics to secondary organs and tissues, including the central nervous system (CNS) (1, 2). Reoviruses primarily disseminate via the blood, although serotype 3 (T3) reoviruses also spread by neural routes (3). In the intestine, reovirus infects intestinal epithelial cells (IECs) and cells in the Peyer’s patch (PP) (4–6). Reovirus is hypothesized to traffic through the mesenteric lymph node (MLN) to the blood via the lymphatics (7). However, the functional route of reovirus systemic spread is not defined.

Reovirus dissemination is influenced by a combination of host and viral factors (8). A key host determinant of reovirus spread is junctional adhesion molecule-A (JAMA), a cell surface receptor for reovirus (9, 10). JAM-A is a tight junction protein that promotes polarization and barrier formation by epithelial and endothelial cells (11, 12). JAM-A also is expressed on monocytes, lymphocytes, dendritic cells, and platelets where it aids in cell migration and extravasation (11, 13, 14). Even though JAM-A is dispensable for reovirus replication in the intestine, it is required for hematogenous spread (10). JAM-A on endothelial cells is required for establishment of viremia, as well as egress of reovirus from the bloodstream into organs (15). A viral factor required for reovirus systemic spread is non-structural protein σ1s (16). Similar to JAM-A, σ1s is dispensable for reovirus replication in the intestine, but is essential for trafficking to the bloodstream (17, 18). The 14 kDa σ1s protein is encoded by the S1 gene segment, which also encodes the attachment protein σ1 (7). Although σ1s is not required for reovirus to traffic from the PP to the MLN, σ1s is necessary for the increase in viral MLN titers observed following oral infection (17). This result suggests that σ1s is necessary for efficient reovirus replication in the MLN. This data further indicates that σ1s is required for viral spread from the PP through intestinal lymphatics to the MLN and ultimately the bloodstream, thereby allowing the establishment of viremia and dissemination to sites of secondary replication. In culture, σ1s facilitates reovirus replication in numerous cell lines, including SV40-immortalized endothelial cells (SVECs) and murine embryonic fibroblasts (MEFs) (16, 19–21). In these cell lines, σ1s functions as a replication accessory factor that promotes reovirus protein synthesis (21). Therefore, σ1s may promote efficient viral replication in cells that are required for reovirus dissemination.

Type-I interferons (IFN-1) are critical for host control of viral infections (22). IFN-1 (IFN-α/β) is produced in responses viral infection and signals in an autocrine or paracrine manner to induce expression of hundreds of IFN stimulated genes (ISGs) that function to limit viral replication through a variety of mechanisms (23–25). Adult mice are normally refractory to reovirus disease. However, reovirus infection is lethal in adult mice that lack interferon-a receptor subunit 1 (IFNAR1) and cannot respond to IFN-1 (26). In IFNAR1^-/-^ mice, IFNAR1 expression on hematopoietic cell is required for protection from reovirus (26). Neonatal mice also succumb more rapidly to reovirus and have higher viral loads than wild-type mice (27, 28). Although IFN-1 is an important host response against reoviruses *in vivo*, the cell types responsible for IFN-1-mediated protection against reoviruses are not defined.

Here, we used tissue-specific IFNAR1 deletion in combination with the IFN-1-sensitive reovirus as a tool to identify a role for lymphatics in reovirus dissemination. We found that σ1s is a viral determinant of reovirus resistance to IFN-1 responses in cultured cells and *in vivo*, as σ1s-deficient reovirus disseminates efficiently in IFNAR1^-/-^ mice. Using Cre-lox technology we found that the IFN-1-sensitive σ1s-deficient reovirus disseminated in mice with lymphatic endothelial cells-specific deletion of IFNAR1. Together, our results indicate that IFN-1 responses in lymphatics are a critical barrier that reovirus must overcome to spread systemically.

## RESULTS

### σ1s facilitates reovirus replication in the presence of IFN-1 responses

To determine the relationship between σ1s and IFN-1 responses, we tested whether σ1s promotes reovirus replication in cells with an intact IFN-1 response. Murine SV40-immortalized endothelial cells (SVEC 4-10; SVECs), a mouse lymphatic endothelial cell line, were infected with T1L or T1L σ1s-null in the presence of isotype control or anti-IFNAR1 antibodies and assessed viral spread in culture over a 7 day period (Fig 1A). T1L produced large foci and spread throughout the culture in the presence of the control antibody. In contrast, T1L σ1s-null was limited to individual cells with no multicellular foci apparent. Treatment with anti-IFNAR1 antibodies led to T1L σ1s-null spread and production of numerous large infected foci. To quantitatively assess the effect of IFN-1 responses on viral replication, we measured viral yields from wild-type and SVECs in which *IFNAR1* was deleted using CRISPR-Cas9 editing (IFNAR1-CRISPR SVECs). Consistent with our previous work (29), T1L produced approximately 10-fold more virus than T1L σ1s-null in wild-type SVECs. In contrast, T1L and T1L σ1s-null replicated to equivalent levels in IFNAR1-CRISPR SVECs (Fig 1B). Similar results were obtained in *IFNAR1^+/+^* and *IFNAR1*^-/-^ MEFs (Fig 1C). T1L replicated to significantly higher levels than T1L σ1s-null in *IFNAR1^+/+^* MEFs, but T1L and T1L σ1s-null produced comparable yields in *IFNAR1^-/-^* MEFs. These results indicate that σ1s facilitates efficient reovirus replication in the presence of IFN-1 responses.

**Figure 1.**
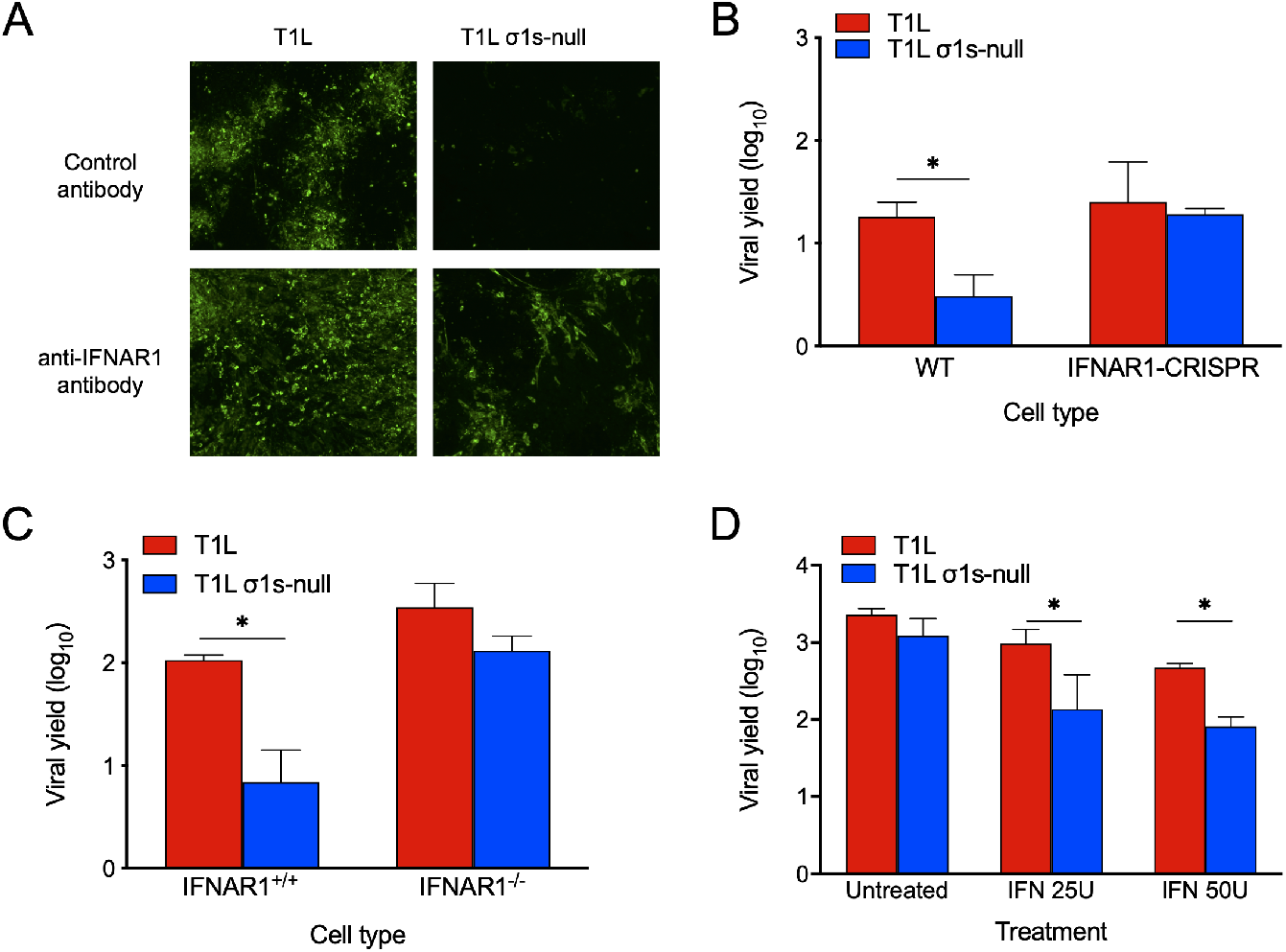
The σ1s protein facilitates reovirus replication in the presence of IFN-1 responses. (A) SVECs were treated with control or anti-IFNAR1 antibodies immediately following infection with T1L or T1L σ1s-null at an MOI of 10 PFU/cell. At 7 days, infected cells were identified via indirect immunofuorescence using reovirus polyclonal antiserum. (B) Wild-type or IFNAR1-CRISPR SVECs or (C) wild-type (*IFNAR1*^+/+^) or *IFNAR1^-/-^* MEFs were infected with T1L or T1L σ1s-null at an MOI of 1 PFU/cell. Viral titers were determined at 0 and 24 h, and results are presented as the mean viral yield from three independent experiments. (D) L929 cells were left untreated or treated with 25 or 50 units of recombinant IFN-α/β for 6 h prior to infection with T1L or T1L σ1s-null at an MOI of 1 PFU/cell. Viral titers were determined at 0 and 24 h, and results are presented as the mean viral yield. Error bars represent SD. *, *p* < 0.05 as determined by Student’s *t*-test.

To determine whether σ1s enhances reovirus sensitivity to IFN-1, replication of T1L and T1L σ1s-null was measured in L929 cells treated with recombinant IFNβ prior to infection (Fig 1D) (30). Unlike SVECs or MEFs, σ1s is not required for reovirus replication in L929 cells and allows assessment of the relationship between σ1s and IFN-1 independent of σ1s effects on viral replication (21, 31). Consistent with previous work, T1L and T1L σ1s-null replicated equivalently in untreated L929 cells. Whereas T1L replication was modestly impaired by IFNβ, the reduction in T1L σ1s-null yields were significantly more pronounced. Together, these findings indicate that σ1s enhances reovirus resistance to IFN-1.

The σ1s protein enhances reovirus replication in SVECs and MEFs by promoting viral protein production (29). Inhibition of viral protein synthesis is a key mechanism by which IFN-1 responses combat viral infections (22). To determine whether σ1s allows reovirus to overcome IFN-1-mediated protein synthesis inhibition, viral protein production was assessed in wild-type and IFNAR1-CRISPR SVECs (Fig 2A) and *IFNAR1^+/+^* and *IFNAR1^-/-^* MEFs (Fig 2B). Consistent with previous results (21), T1L produced more viral protein than T1L σ1s-null in wild-type SVECs and MEFs. Although both viruses produced more protein in IFNAR1-CRISPR SVECs and *IFNAR1^-/-^* MEFs than in wild-type cells, protein expression by T1L σ1s-null remained substantially lower than T1L in both cell types. Together, these data indicate that σ1s does not directly counteract the inhibition of reovirus protein synthesis caused by IFN-1.

**Figure 2.**
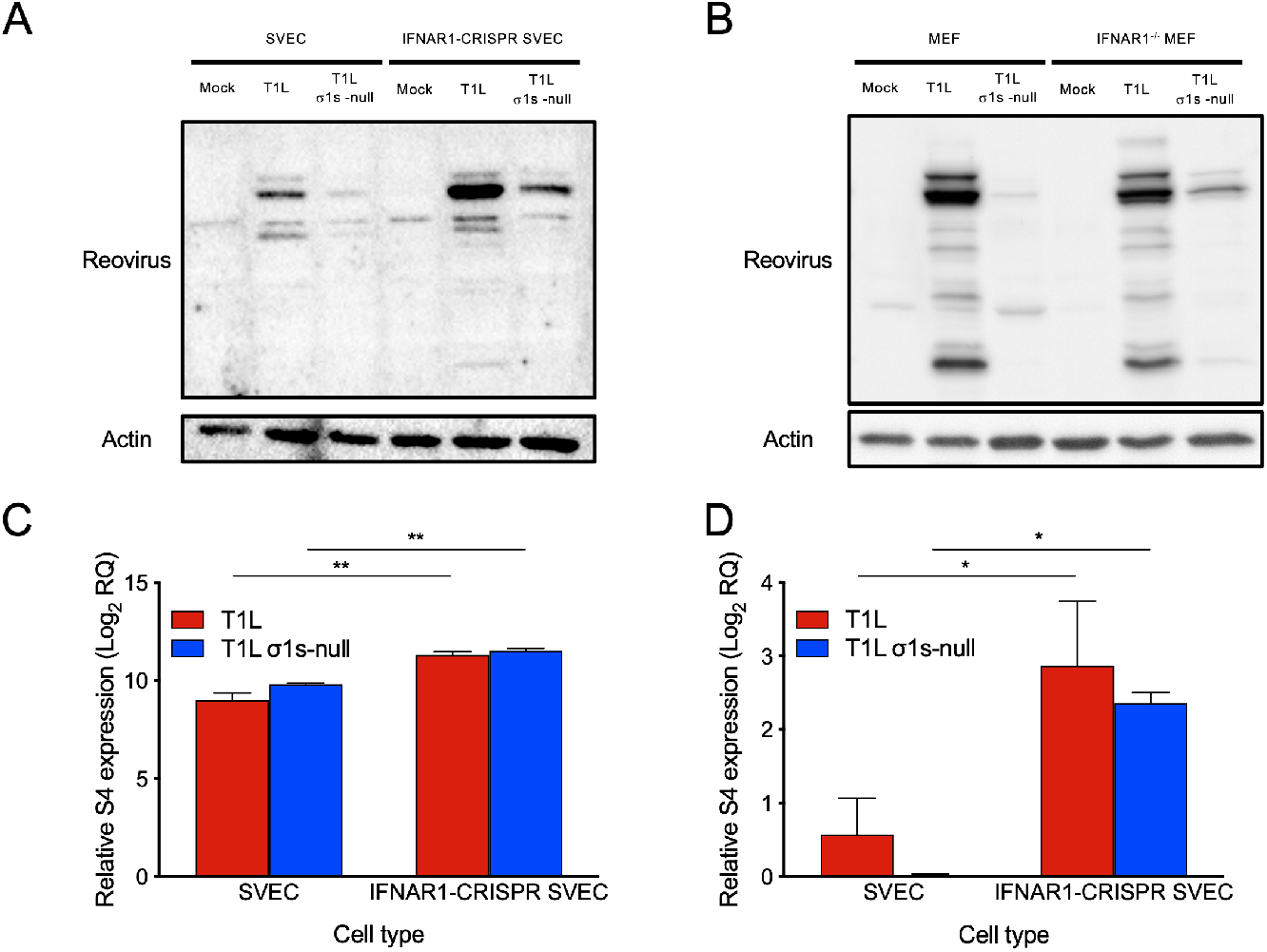
The σ1s protein does not counterract IFN-1-mediated inhbition of reovirus protein synthesis. (A) Wild-type or IFNAR1-CRISPR SVECs or (B) wild-type (*IFNAR1^+/+^*) or *IFNAR1*^-/-^ MEFs were infected with T1L or T1L σ1s-null at an MOI of 10 PFU/cell. At 18 h, whole cell lysates were collected and seperated by SDS-PAGE. Reovirus proteins were detected by western blot. β-actin was detected as a loading control. (C and D) WT or IFNAR1-CRISPR SVECs were infected with T1L or T1L σ1s-null at an MOI of 10 PFU/cell. At 0 or 18 h, total RNA was collected and relative expression of (C) mRNA or (D) negative-sense RNA was determined compared to 0 h. The RQ of positive- or negative-sense RNA was determined with reference to the quantity at the 0 h. The RQ of reovirus mRNA was determined by subtracting the RQ of negative-sense viral RNA (representing genomic RNA production) from the RQ of positive-sense RNA. Data are presented as the mean log2 RQ from three independent experiments. Error bars indicate standard deviations. *, *p* < 0.05; **, *p* < 0.005 as determined by Student’s *t*-test.

We next quantified viral RNA to determine the effect of IFN-1 responses on viral RNA synthesis by T1L and T1L σ1s-null. Consistent with previous results (21), viral S4 mRNA (Fig 2C) and negative-sense RNA (Fig 2D) levels were similar between T1L and T1L σ1s-null in wild-type SVECs. T1L and T1L σ1s-null produced approximately four-fold more S4 mRNA and negative-sense RNA in IFNAR1-CRISPR SVECs compared to control cells. Like control cells, S4 mRNA and negative-sense RNA levels were comparable between T1L and T1L σ1s-null. These data are consistent with previous findings that σ1s is dispensable for reovirus RNA synthesis (29). These results indicate that while IFN-1 responses limit reovirus RNA synthesis, σ1s does not specifically modulate antiviral responses that prevent viral RNA accumulation. These results further suggest that T1L σ1s-null replication in cells lacking IFN-1 responses may result from enhanced viral RNA synthesis, which increases viral protein synthesis and progeny production.

### The σ1s protein facilitates reovirus dissemination in the face of IFN-1 responses *in vivo*

Our data and published studies (32) indicate that σ1s contributes to reovirus IFN-1 resistance in cultured cells. To determine whether σ1s is required for reovirus to overcome IFN-1 responses *in vivo*, we assessed T1L and T1L σ1s-null spread in wild-type and *IFNAR1*^-/-^ mice (Fig 3). T1L replicated in the intestine and spread systemically in wild-type mice. Consistent with previous findings (16), although T1L σ1s-null produced comparable viral titers to T1L in the intestine of wild-type mice, T1L σ1s-null titers in peripheral organs (brain, heart, liver, and spleen) were near or below the limit of detection. As in wild-type mice, T1L and T1L σ1s-null produced comparable intestinal titers in *IFNAR1*^-/-^ mice. However, equivalent titers of T1L and T1L σ1s-null were recovered from peripheral organs of *IFNAR1*^-/-^ mice. These data indicate that the IFN-1 response acts as a barrier to reovirus hematogenous dissemination. These findings further suggest that σ1s is required for reovirus spread systemically in the presence of IFN-1 responses.

**Figure 3.**
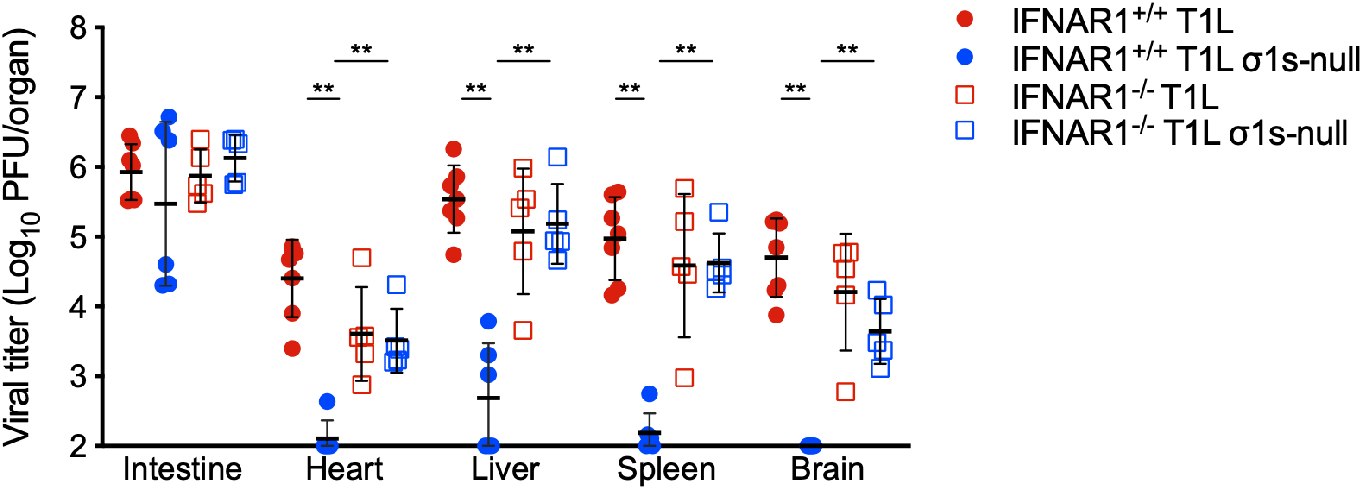
The σ1s protein is required for reovirus dissemination in the presence of IFN-1 responses. Three-to-four day-old wild-type (*IFNAR1^+/+^*) or *IFNAR1*^-/-^ neonatal mice were infected orally with 10^4^ PFU of T1L or T1L σ1s-null. At 4 days, the indicated organs were resected, homogenized, and viral titer was determined by plaque assay. Error bars represent SD. **, *p* < 0.005 as determined by Mann-Whitney test.

### The σ1s protein is required for efficient reovirus replication in lymphatic endothelial cells

Reovirus is hypothesized to spread via the lymphatics, which are largely formed from lymphatic endothelial cells (LECs) (33). Previous work revealed that σ1s was required for efficient reovirus replication in SVECs, an immortalized lymphatic endothelial cell line (21). To determine whether σ1s is required for reovirus replication in primary LECs, we quantified T1L and T1L σ1s-null progeny yields produced by LECs derived from C57BL/6 mice (Fig 4A). We found that T1L generated significantly higher progeny yields than T1L σ1s-null in primary LECs at both MOIs tested. In cells where σ1s is required for reovirus replication, σ1s also mediates optimal viral protein production (21). In primary LECs, we observed that differential replication of wild-type and σ1s-deficient viruses correlated with differences in viral protein production, as T1L produced substantially more viral protein than T1L σ1s-null (Fig 4B). Together, these data indicate that σ1s promotes reovirus replication in primary LECs by enhancing viral protein production.

**Figure 4.**
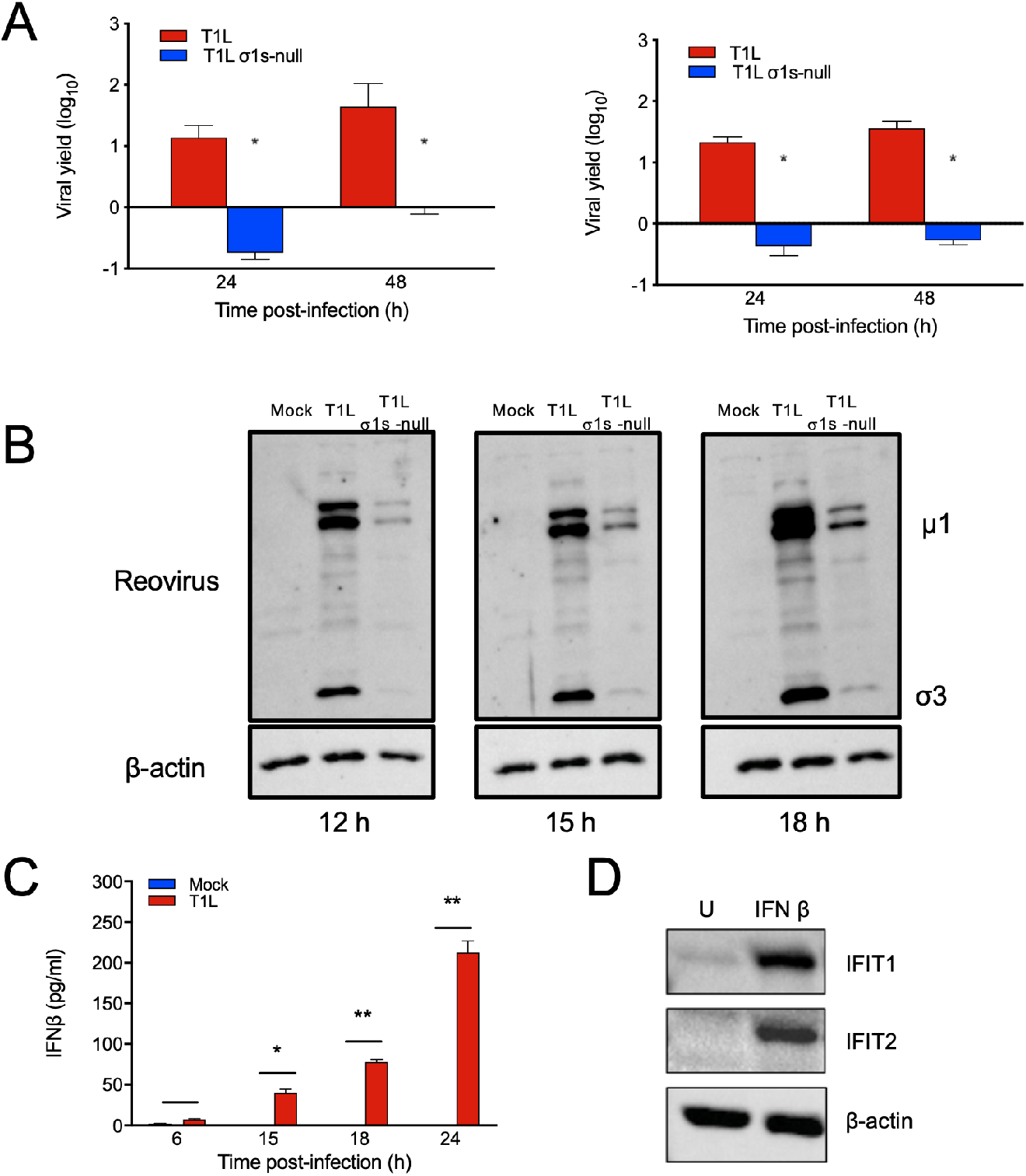
The σ1s protein facilitates reovirus protein synthesis and replication in primary lymphatic endothelial cells. (A) Primary LECs from C57BL/6 mice were infected with T1L or T1L σ1s-null at MOIs of 1 (left) or 10 (right) PFU/cell. At 0, 24, and 48 h, viral titers were determined by plaque assay. Results are presented as the mean viral yield from two independent experiments. Error bars represent SD. *, p < 0.05 as determined by Student’s *t*-test. (B) Primary LECs were mock infected or infected with T1L or T1L at an MOI of 5 PFU/cell. At the indicated times, whole cell lysates were collected and seperated by SDS-PAGE. Reovirus proteins were detected by western blot. β-actin was detected as a loading control. (C) Primary LECs were mock infected or infected with T1L or T1L σ1s-null at an MOI of 100 PFU/cell. At the indicated times, IFNβ levels in the supernatants were determined by ELISA. (D) Primary LECs were treated with 200 U IFNβ or left untreated. At 6 h, whole cell lysates were collected and seperated by SDS-PAGE. IFIT1 and IFIT2 were detected by western blot. β-actin was detected as a loading control.

### Lymphatics facilitate reovirus dissemination

If lymphatics function as conduits for reovirus dissemination, we hypothesized that ablating IFN-1 responses specifically in lymphatic endothelial cells would enable dissemination of σ1s-deficient reovirus. To test this hypothesis, we used the IFN-1-sensitive σ1s-null reovirus in combination with lymphatic-specific deletion of *IFNAR1.* We first confirmed that primary LECs secrete IFNβ in response to reovirus infection (Fig 4C) and produce ISGs following IFN-1 treatment (Fig 4D). To generate lymphatic-specific *IFNAR1* deletion mice, *IFNAR1^fl/fl^* mice were crossed with *Lyve1-Cre* mice (34, 35). LYVE-1 is a receptor for hyaluronan that promotes LEC proliferation (36, 37). LYVE-1 is a commonly used marker for the lymphatic endothelium, but also is expressed on liver sinusoid, some tissue-resident macrophages, and a subset of hematopoietic stem cells (34, 38–40). T1L disseminated in the parental *IFNAR1^fl/fl^* (Fig 5A) and *Lyve1-Cre* (Fig 5B) strains following oral inoculation, similar to results obtained with C57BL/6 mice (18). In contrast, T1L σ1s-null did not spread efficiently in either parental mouse strain.

**Figure 5.**
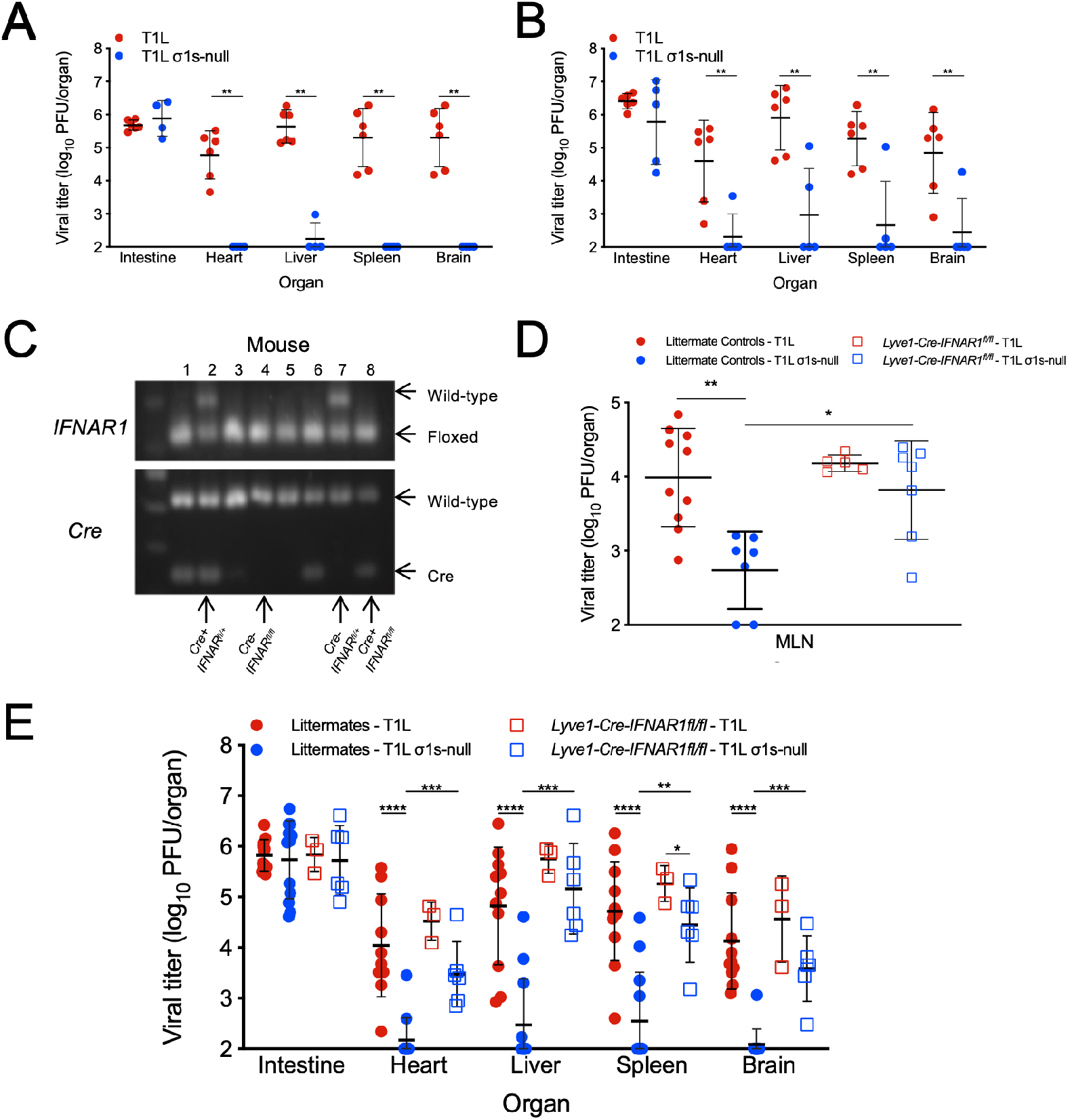
IFN-1 responses in lymphatics limits reovirus dissemination. (A) *IFNAR1^fl/fl^* or (B) *Lyve1-Cre* mice were orally infected with 10^4^ PFU T1L or T1L σ1s-null. At 4 days, the indicated organs were resected, homogenized, and viral titer was determined by plaque assay. (C) The genotype of transgenic mice was determined by performing PCR analysis of chromosomal DNA using primers specific for IFNAR1 and Cre. The floxed IFNAR1 alleles and Cre were differentiated based on migration of the PCR products in 1% agarose gel and stained with EtBr. (D and E) Littermate control (*IFNAR1^fl/fl^, IFNAR1^+/fl^, Lyve1-Cre-IFNAR1^+/fl^*) or *Lyve1-Cre-IFNAR1^fl/fl^* mice were orally infected with 10^4^ PFU T1L or T1L σ1 s-null. At 4 days, (D) MLNs or (E) the indicated organs were resected, homogenized, and viral titer was determined by plaque assay. Error bars represent SD. *, *p* < 0.05; **, *p*<0.01; ***, *p* < 0.0005; ****, *p*<0.0001 as determined Mann-Whitney test.

The F1 progeny resulting from crossing *IFNAR1^fl/fl^* and *Lyve1-Cre* strains were bred to *IFNAR1^fl/fl^* mice. The resulting progeny *(Lyve1-Cre-IFNAR1^+/fl^, IFNAR1^+/fl^, IFNAR1^fl/fl^* littermate controls and *Lyve1-Cre-IFNAR1^fl/fl^* LEC IFNAR1 deletions) were infected with T1L or T1L σ1s-null and at 4 days post-infection viral tissue titers were determined. Mice were genotyped by PCR at the time of harvest to determine their *Cre* and *IFNAR1* status (Fig 5C). Following oral infection, we found that T1L titers were substantially higher on day 4 than T1L σ1s-null in the MLNs of littermate control animals (Fig 5D). In contrast, T1L and T1L σ1s-null produced comparable titers in MLNs from *Lyve1-Cre-IFNAR1^fl/fl^* mice. These data indicate that IFN-1 responses in the lymphatics impair spread of σ1s-deficient reovirus to the MLN.

We next assessed reovirus spread in mice with lymphatic *IFNAR1* deletion. In littermate control mice, T1L and T1L σ1s-null exhibited comparable titers in the intestine, but only T1L had high titers in peripheral organs. T1L σ1s-null titers in organs from littermate control mice were near or below the level of detection (Fig 5E). These data are consistent with observations in wild-type, *IFNAR1^fl/fl^*, and *Lyve1-Cre* mice that σ1s is required for efficient reovirus dissemination. In contrast, T1L and T1L σ1s-null produced largely comparable titers in all organs of *Lyve1-Cre-IFNAR1^fl/fl^* mice. Thus, deletion of *IFNAR1* in lymphatics enables dissemination of σ1s-deficient reovirus. Taken together, these results indicate that lymphatic IFN-1 responses are critical for controlling reovirus dissemination.

## DISCUSSION

Here, we identified a role for lymphatic IFN-1 responses in controlling hematogenous reovirus dissemination. In the intestine, reovirus is transcytosed by microfold cells (M-cells) in the gut-associated lymphoid tissue (GALT) where it infects the basolateral surface of IECs (8). Replication in IECs mediates reovirus release into the stool for dissemination to future hosts (8, 41, 42). To disseminate, reovirus is taken up by cells in the Peyer’s patch (4, 10, 18) and hypothesized to traffic through the MLN via the lymphatics and then to the bloodstream (8). However, the operant route of reovirus dissemination is not defined. Our data provide support for the hypothesis that the lymphatics function in hematogenous reovirus dissemination, as IFNAR1 deletion in LYVE-1-expressing cells allowed spread of the IFN-1-senstive σ1s-deficient reovirus.

Global and conditional deletion of JAM-A revealed that endothelial cells, but not hematopoietic cells, mediate establishment of reovirus viremia and egress from the blood into tissues (10, 43). It is possible that σ1s promotes reovirus replication in LECs that line lymphatic vessels and lymph nodes to provide a reservoir that seeds virus for trafficking through the lymphatics to the blood. Consistent with this hypothesis, we found that σ1s was required for reovirus protein expression and replication in primary LECs (Fig 4). LYVE-1 is predominantly expressed on LECs, but also reported to be expressed on a small subset of fetal and adult hematopoietic stem cells, liver blood sinusoids, and adult tissue-resident macrophages (34, 38–40). Conditional expression of JAM-A on hematopoietic cells is insufficient to restore reovirus hematogenous spread in JAM-A-deficient mice, indicating that hematopoietic cells do not mediate reovirus dissemination (43). It also is unlikely that reovirus accesses liver sinusoids following oral inoculation. Finally, our studies use neonatal mice and adult tissue-resident macrophages would not be involved in dissemination.

It also is possible that loss of IFN-1 signaling in LYVE-1-positive cells increases the permeability of the lymphatic endothelium, thereby allowing reovirus to escape the lymphatic vessels. IFN-1 controls LEC expansion in response to viral infection (44) and also modulates vascular endothelial barrier function, particularly at the blood-brain barrier (45–47). Lack of IFN-1 signaling in the lymphatic endothelium could allow the σ1s-null virus to leak into the lymphatic vessels and traffic to the blood. Finally, loss of IFN-1 responses in the LECs could alter the transport dynamics of the lymphatics. In the skin, IFN-1 signaling blocks fluid transport to the regional lymph node and limits poxvirus dissemination (48). If IFN-1 has similar effects on the dynamics of gut lymphatics, removing IFNAR1 from LECs could alleviate the normal interruption of lymphatic flow designed to impede viral dissemination.

The σ1s protein is dispensable for reovirus replication in the intestine, but is required for dissemination to peripheral organs (18). Our data suggest that the IFN-1 system restricts reovirus spread in the absence of σ1s. However, why σ1s is dispensable for reovirus replication in the intestine remains an open question. One possibility is that reovirus replication in the intestine is largely controlled by interferon-λ (IFN-λ), as opposed to IFN-1 (41, 42, 49). Expression of the IFN-λ receptor, IFN-λ receptor 1 (IFNLR1), is largely restricted to the epithelial compartment, which also lacks IFNAR expression (50, 51). Consequently, IFN-λ protects IECs whereas IFN-1 functions in non-epithelial tissues (52). Like IFN-1, IFN-λ induces ISG expression, but IFN-1 induces ISG with more rapid kinetics than IFN-λ and the magnitude of the antiviral response is greater for IFN-1 than IFN-λ (53). IFN-λ limits reovirus replication in the intestine, as mice lacking IFNLR1 or IFN-λ2/3 have elevated reovirus IEC infection and shedding (41, 42, 54). We found no difference in viral intestinal titers between wild-type and σ1s-deficient viruses in wild-type or IFNAR1-knockout mice. These data are consistent with IFN-λ as the primary means of controlling reovirus replication in the intestine. These findings also suggest that σ1s is not required for reovirus to overcome IFN-λ. It is possible that σ1s is more important for resisting IFN-1 than IFN-λ responses due to the lower potency of IFN-λ compared to IFN-1. However, the relationship between σ1s and IFN-λ remains to be explored.

Like most viruses, reovirus activates a variety of cellular mechanisms that function to impair viral protein synthesis, including PKR, which phosphorylates eIF2α to block translation (55–57) and the OAS-RNaseL system that mediates RNA degradation (55). Reovirus must produce viral proteins in the face of host translational shutoff in order to replicate efficiently. Reovirus may even benefit from host shutoff, as viral replication is decreased in MEFs lacking PKR or expressing a constitutively active form of eIF2α (57). Reovirus uses multiple mechanisms to evade host translational shutoff, including outer capsid protein σ3 binding dsRNA to blunt PKR activation (58) and non-structural protein σNS facilitating escape from host shutoff by mediating dissolution of stress granules (59). However, reovirus has other means to circumvent host shutoff responses. It is possible that σ1s promotes reovirus protein expression by counteracting the function of one or more ISGs that block host translation (22, 60). Although σ1s is required for reovirus replication in the presence of IFN-1 responses, σ1s does not function as a classical IFN-1 antagonist (21), as IFN-1 secretion, IFNAR signaling, and ISG induction are comparable between wild-type and σ1s-deficient viruses (21). We observed that viral protein expression by the σ1s-deficient virus is only partially restored in the absence of IFNAR1 signaling. This result suggests that σ1s promotes reovirus protein expression via an IFN-1-independent mechanism.

The σ1s protein is required for systemic reovirus spread (17, 18). Here, we found that σ1s is important for reovirus resistance to IFN-1 in cell culture and *in vivo.* We used the IFN-1-sensitive σ1s-deficient reovirus in combination with tissue-specific deletion of IFNAR1 in lymphatic endothelial cells to identify a role for IFN-1 responses in lymphatics in controlling reovirus spread. Together, our findings provide new insight into mechanisms that control reovirus dissemination and further define how reovirus spreads from mucosal sites of infection to target organs and tissues.

## MATERIALS AND METHODS

### Cells and viruses

Murine L929 fibroblasts were maintained in Joklik’s modified Eagle medium (JMEM, Sigma) supplemented with 10% heat-inactivated fetal bovine serum (FBS, Invitrogen), 1% L-glutamine (Invitrogen), 1% penicillin/streptomycin (Invitrogen), and 0.1% amphotericin B (Sigma). SV-40 immortalized endothelial cells (SVECs, ATCC), C57BL/6 murine embryonic fibroblasts (MEFs), IFNAR1^-/-^ MEFs, and human embryonic kidney 293 cells (HEK293T) were maintained in Dulbecco’s modified Eagle medium (DMEM, Invitrogen) supplemented to contain 10% heat-inactivated FBS and 1% L-glutamine. SVECs lacking the IFN-α/β receptor (IFNAR1-CRISPR) were maintained in the same media as SVECs with 2 μg/mL puromycin (Gibco). Primary LECs (Cell Biologics) were maintained in Endothelial Cell Medium (Cell Biologics) supplemented to contain 5% FBS, 0.1% VEGF, 0.1% heparin, 0.1% endothelial cell growth supplement (ECGS), 0.1% epidermal growth factor (EGF), 0.1% hydrocortisone, 1% L-glutamine, and 1% antibiotic-antimycotic solution.

Reoviruses were generated using plasmid-based reverse genetics as described previously (8, 16, 19, 61, 62). Purified reovirus stocks were obtained from second- or third-passage L929 cell lysates from twice-plaque-purified reovirus (63). Vertrel was used to extract reovirus particles, which were separated on a 1.2-1.4 g/cm^3^ CsCl density gradient and exhaustively dialyzed in virion storage buffer (150 mM NaCl, 15 mM MgCl_2_, 10 mM Tris-HCl pH 7.8). Reovirus stocks were titered by plaque assay on L929 fibroblasts (64).

### CRISPR-Cas9 deletion of IFNAR1

IFNAR1 was deleted in SVECs using the CRISPR-Cas9 system (65). The plentiCRISPRV.2 (Addgene) was digested with *BsmBI* and ligated with guide RNA sequences specific for IFNAR1, (IFNAR1 +) 5’-CACCGGCTCGCTGTCTGTGGCGCGG-3’ and (IFNAR1-) 5’-AAACCCGCGCCCACGACAGCGAGCC-3’. The cloned plasmids were transfected in to HEK293T cells in combination with pSPAX2 and pCMV-G plasmids using Lipofectemine 2000 (Invitrogen). Supernatants were collected at 24 and 48 h post-transfection, passed through a 0.45 μm syringe filters, and applied to SVECs in 6-well plates (~50% confluent). At 48 h post-transduction, puromycin (Invitrogen, 2 μg/mL) was added. Puromycin-selected SVECs were tested for IFNAR1 deletion by treatment with IFN-β (PBL) followed by RT-qPCR to measure ISG expression.

### Viral replication assays

Monolayers of cells in 24 well plates (1×10^5^ cells/well) were infected with T1L or T1L σ1s-null at an MOI of 1 PFU/cell at 4°C for 1 h. After adsorption, cells were washed twice with cold PBS and fresh media was added. Infected cells were freeze-thawed twice at the indicated times in the figure legends and viral titer was determined by plaque assay on L929 fibroblasts (64). Viral yields were calculated using the formula: log_10_yield_tx_ = log_10_(PFU/mL)_tx_ – log_10_(PFU/mL)_t0_ where tx is the time post-infection.

### Immunoblotting

Monolayers of cells in 6-well plates (1×10^6^ cells/well) were mock infected or infected reovirus or treated with recombinant IFNβ (PBL Assay Science) as indicated in the figure legends. Whole cell lysates were collected in RIPA buffer (20 mM Tris pH 7.4, 150 mM NaCl, 1mM EDTA, 1% sodium dodecyl sulfate, 1% desoxycholate, 1/100 IGEPAL [NP-40]) at the indicated times. Total protein in each sample was quantified using the DC protein assay (BioRad) and 25 μg protein were separated by SDS-PAGE. Proteins were transferred to nitrocellulose membrane and incubated in blocking buffer (5% milk in 1X TBS with 0.05% Tween-20 [TBS-T]) for 1 h. Membranes were incubated in blocking buffer containing reovirus-specific rabbit polyclonal anti-serum (1:2000 dilution), (need info for IFIT1 and IFIT2 antibodies) overnight at 4°C. Membranes were washed three times with TBS-T followed by incubation in blocking buffer containing horseradish peroxidase-conjugated goat anti-rabbit IgG (Jackson Immunolabs, 1:2000 dilution) for 1 h with rocking. Following three TBS-T washes, proteins were detected using SuperSignal West Chemiluminescent Substrate (ThermoFisher). Blots were stripped for re-probing by washing membranes three times with TBS followed by incubation in Restore Western Blot Stripping buffer (Thermo Scientific) for 15 min at RT. Following three washes in TBS, membranes were blocked as described above and β-actin was detected using mouse β-acting specific monoclonal antibody (Sigma, 1:10,000 dilution) and peroxidase-conjugated goat anti-mouse IgG (Jackson Immunolabs, 1:2000 dilution).

### RT-qPCR

Monolayers of cells in 6 well plates (1×10^6^ cells/well) were infected with T1L or T1L σ1s-null at an MOI of 10 PFU/cell. Total RNA was collected using the RNeasy Plus kit (QIAGEN). Reovirus S4 RNA was quantified using the Taq-Man Fast Virus One-Step Master Mix (Applied Biosystems) and GAPDH was detected as an endogenous control using the Pre-Developed TaqMan Assay Reagent kit for mouse GAPDH (Applied Biosystems) as described previously (21). The relative quantity (RQ) of reovirus positive- or negative-sense RNA was quantified using t = 0 h post-infection as the reference sample. the ΔΔC_T_ was calculated for each sample by the formula:

ΔΔC_T_ = (Unknown CT_tx_ – GAPDH CT_tx_) – (Unknown CT_t0_ –GAPDH CT_t0_) where tx = time post-infection. The ΔΔCT was then used to calculate RQ by the formula: RQ = 2^Λ-ΔΔC_T_^.

### Mouse experiments

Animal husbandry, housing, and experiments were performed according to the guidelines of the Division of Laboratory Animal Medicine (DLAM) at UAMS. C57BL/6 (JAX stock #000664), C57BL/6 *IFNAR1*^-/-^ (JAX stock #028288), C57BL/6 *IFNAR1^fl/fl^* (JAX stock #028256), and C57BL/6 *Lyve1-Cre* (JAX stock #012601) mice were obtained from Jackson Laboratory. *IFNAR1^fl/fl^* and *Lyve1-Cre* mice were crossed to obtain *IFNAR1^+/fl^/Lyve1-Cre* heterozygous mice. *IFNAR1^+/fl^/LYVE1-Cre* heterozygous mice were crossed to *IFNAR1^fl/fl^* mice for experiments, yielding litters of *IFNAR1^fl/fl^, IFNAR1^+/fl^, IFNAR1^+/fl^/Lyve1-Cre*, and *IFNAR1^fl/fl^/Lyve1-Cre* mice. Three-four day old mice were infected orally with 10^4^ PFU T1L or T1L σ1s-null diluted in PBS as previously described (10, 66). At 4 days post-infection, organs were resected, homogenized, and viral titer was determined by plaque assay. Infected mice were genotyped after experiments using the KAPA HotStart Mouse Genotyping Kit (KAPA Biosystems) and primers for the floxed *IFNAR1* allele and the *Lyve1-Cre* gene from the Jackson Laboratory website.

### Statistics

Differences in viral replication were determined by an unpaired Student’s *t*-test. Differences in viral titer from mouse experiments were determined by Mann-Whitney test. Statistical tests were performed using Prism software (GraphPad Software, Inc.). *p* values < 0.05 were considered significant.

## ACKNOWLEDGEMENTS

M.B.P. and K.W.B. conceived and designed the experiments. M.B.P., M.D.Z., M.A.H. and K.W.B. performed the experiments. M.B.P., M.D.Z., and K.W.B. analyzed data. T.W. provided input on experimental design. M.B.P. and K.W.B. drafted the manuscript with critical revision from all other authors. We also thank Pranav Danthi and Craig Forrest for critical reading of the manuscript.

This research was supported by Public Health Award K22 AI90497 (K.W.B.) and R01 AI118801 (K.W.B.). Additional support was provided by the Center for Microbial Pathogenesis and Host Inflammatory Response (P20 GM103625). The pMSCV-puro plasmid and EcoPak cells were provided by Craig Forrest (UAMS).

